# Reducing *Aedes albopictus* breeding sites through education: a study in urban area

**DOI:** 10.1101/385872

**Authors:** A. Stefopoulou, G. Balatsos, A. Petraki, Shannon L. LaDeau, D. Papachristos, A. Michaelakis

**Affiliations:** Benaki Phytopathological Institute, Department of Entomology and Agricultural Zoology, 14561, Kifissia.; Cary Institute of Ecosystem Studies, Millbrook, New York, United States of America

## Abstract

*Aedes albopictus* tends to proliferate in small, often man-made bodies of water, largely present in urban private areas. For this reason, education and community participation are considered crucial for source reduction and mosquito control. In the current study, we tried to relate for the first time in Greece, the effectiveness of resident education in an urban area with the number of breeding sites based on previous KAP (knowledge, attitudes and practices) studies. Our study examines the relationship between mosquito breeding sources and socioeconomic or demographic characteristics of different households in a Greek municipality and evaluates efficacy of resident education. The results revealed that only a minority of residents knew where mosquitoes breed (18.6%) and only 46% felt that residents had any responsibility for managing breeding habitat. Our findings strongly suggest that only the presence of scientific staff inspecting possible habitats in their properties, could be enough to stimulate practices towards source reduction. However, educational interventions alone with printed education material cannot enhance significant community participation and source reduction.

## Introduction

The Asian tiger mosquito, *Aedes* (*Stegomyia*) *albopictus* (Skuse, 1895) (Diptera: Culicidae), is considered one of the 100 most invasive species in the world presenting a significant expansion in many parts of the globe [1–3]. *Aedes albopictus* is a vector of viruses for dengue, chikungunya, yellow fever, Japanese encephalitis [4–5], and zoonoses, such as dirofilarioses [6] presenting thus a notable threat to public health. Apart from vector of viruses *Ae. albopictus* causes serious nuisance to citizens.

In Europe, *Ae. albopictus* was first detected in Albania in 1979 [7], while in Greece it was first detected in Thesprotia and Corfu in 2003 [8]. Since then, *Ae. albopictus* has invaded most of the continental part of the country with the exception of some areas in northern Greece and the Aegean islands [3].

*Aedes albopictus* is a container-inhabiting mosquito species and in urban areas can be found in both public and private areas with vegetation [9]. In Greece both Regional Units and municipalities are responsible for mosquito control in public areas (eg catch basins, rivers, streams etc), however urban areas are comprised of a large number of private areas with numerous water containers that work as breeding sites for the Asian tiger mosquito. Given the fact that private areas are inaccessible to prefecture’s control measures and intense chemical control efforts are of low efficacy, resident-based management (i.e. breeding source reduction) is considered a most effective and affordable mean for controlling of mosquitoes [10–11].

Education campaigns and community participation is considered beneficial in the reduction of container habitats and vector control [12] and is recommended by the World Health Organization [13]. In many cases, educational programs have indeed resulted in an important impact in container habitat reduction [12,14]. Education campaigns focus on increasing public participation and awareness on source reduction, however it remains uncertain whether this education is actually accompanied by mosquito population reduction [15–17,11].

For urban mosquito species, such as *Ae. albopictus* and *Cx. pipiens*, the knowledge, attitudes and practices (KAP) of residents can influence the efficacy of management [18]. In the current study, we tried to relate for the first time in Greece, the effectiveness of resident education in an urban area with the number of breeding sites based on previous KAP studies [18, 11]. Our study examines the relationship between mosquito abundance and socioeconomic or demographic characteristics of different households in Greece and evaluates efficacy of resident education. The quantitative KAP survey included predefined standardized questions aiming at revealing misconceptions that could limit the effectiveness of mosquito control practices. Therefore, our KAP study tried to address the following research questions: (a) Are there demographic or housing variables that are associated with greater numbers of existing mosquito habitat at the initial survey period? (b) Is the initial container habitat abundance associated with reported resident knowledge, attitude or practice metrics? (c) Could demographic, KAP, or housing information predict relative container reductions? (d) Did an intervention in the form of resident education result in greater habitat reduction?

## Materials and Methods

### Study site and education materials

Our study employed a KAP questionnaire, household surveys, and introduced an education campaign to evaluate changes in the abundance of breeding sites among the households of the respondents. Preliminary monitoring studies showed the presence of established population of *Ae. albopictus* in all areas of the municipality. The study site was the municipality of Palaio Faliro which is situated 6 km southwest of Athens city Centre, on the east coast of the Saronic Gulf. It has an area of 4.574 km^2^ with almost 64 thousand inhabitants (in 2011, the last census). The Pikrodafni stream flows into sea on the border of Palaio Faliro and Alimos (Image 1).

**Image 1.**
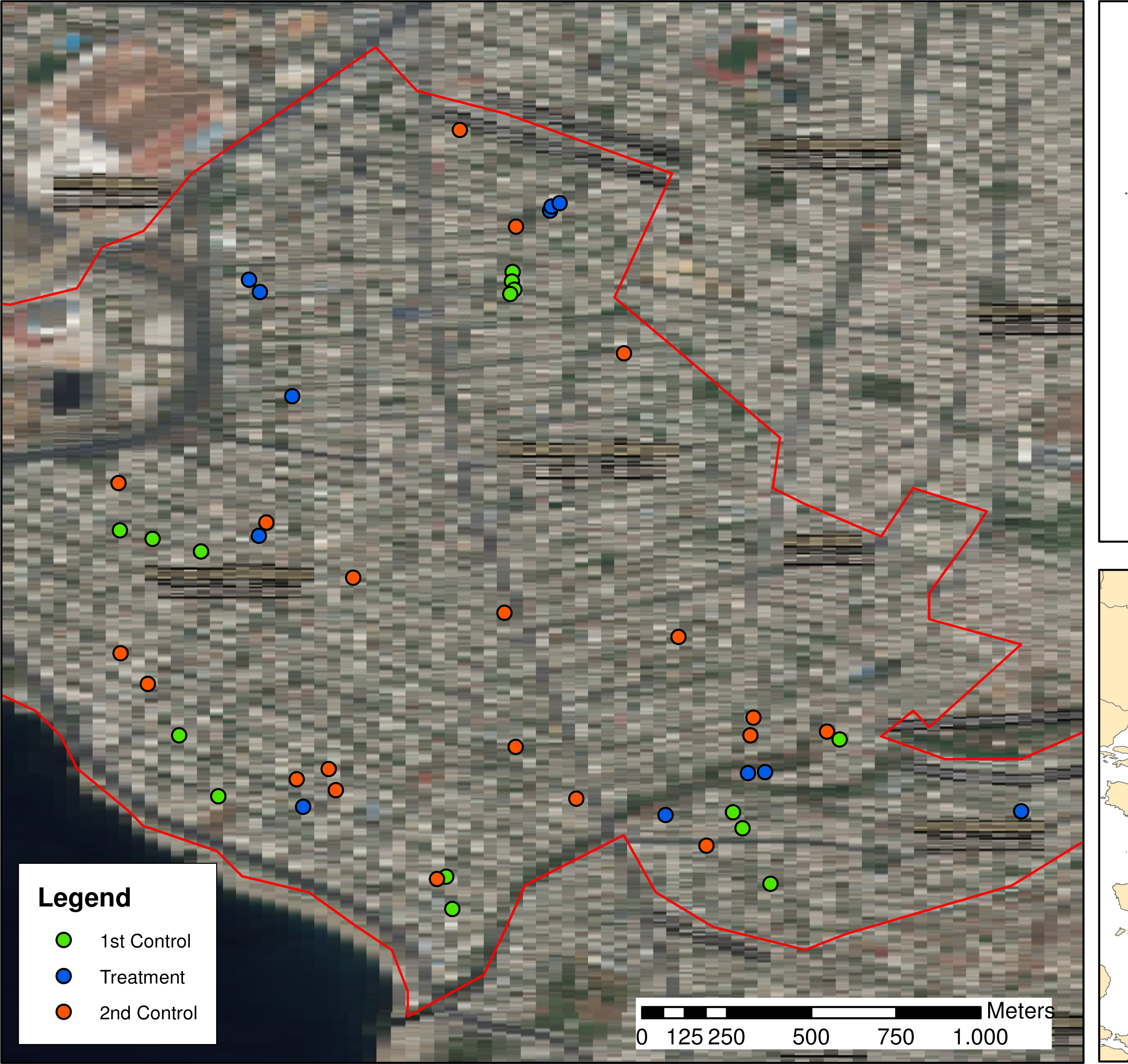
Distribution of under study households in the Municipality of Palaio Faliro (Region of Attica-Greece; see more details for control and treatment households in text)

Since the peak period for *Ae. albopictus* in Athens is typically September-October [19] the first visit in all households was in July 2017 and the second in November 2017. In more details, the first step was to identify the households to be included in our study which were distributed in several neighborhoods/areas of the municipality. We were able to find, with the assistance of the Environmental Department (Environment Unit and the Green) of the Municipality of P. Faliro, 37 different households. During the initial visit to each household (July 2017) researchers recorded the total number of breeding sites that could potentially hold water in each yard. Breeding sites were recorded either as yard (containers put out purposefully for recreation/aesthetic purposes), structural (habitats created by permanent features of the yard or building) or trash. After the first visit, we distributed to 17 randomly selected houses the educational material (Treatment group) while the remaining 20 households were considered as a control group. On November 2017 we revisited all households, both treatment and control group, to record the number of breeding sites. As this is the first assessment with the distribution of education materials and raising awareness in Greece, in the last visit, we also included a new set of 27 households (a 2^nd^ Control group not included in the first visit) aiming to evaluate behaviors that may be influenced because of our first inspection. All visits were completed during the daylight (usually from 9.00 am until 4.00 pm).

A primary aim of our study was to evaluate the effect of an education intervention on household source reduction. Education materials were distributed after our first yard inspection and included written mosquito information, as well as a cap and T-shirt. The written mosquito information included general information about mosquitoes and their breeding sites. The leaflet with information was created in the framework of the project “LIFE CONOPS” (LIFE12 ENV/GR/000466) funded by the European Commission in the framework of the programme LIFE + Environment Policy and Governance. In the supplementary section the Greek version of the leaflet is available (as distributed) while the English version is on LIFE CONOPS website (www.conops.gr)]. All educational materials were delivered personally to each household (hand-delivered to a resident). Thus, treatment households were visited twice in July, once for the yard inspection when breeding source habitats were counted and a second time to distribute education materials. Control households were only visited for the initial yard inspection.

### KAP Questionnaires

At the time of the initial yard inspection we distributed questionnaires to each household (64 in total) to collect information on baseline resident knowledge, attitude and practices (KAP, following similar methodologies in [11,18]). Moreover, the questionnaire was designed to collect demographic information: age, gender, education, household income, ownership and presence of children under 18 years of age. In our study, we included private properties that were apartment complexes (block of flats) and therefore in demographic information we collected data relative to the housing information: if the household was in a complex consisting of less or more than 4 apartments (kind of house) and if the participant (household) was the building administrator for the block of flat. All questionnaires were completed only by adults at each household (>18 years-old). All data was analyzed anonymously.

Following KAP methodology, an overall score was assigned for each respondent concerning their knowledge and motivation related to mosquito control.

#### 1. Knowledge score

The respondents’ knowledge score was based on three questions that aimed at detecting background knowledge about mosquitoes. The first question asked if they know where mosquitoes lay their eggs and grow. Respondents who did not know the answer scored 0. All correct answers scored 1 point. The second question asked them to identify who is responsible for mosquito control. If the respondents reported “residents” among the responsible, they scored “correctly” and got 1 point, otherwise they scored 0 points. The third question asked the respondents to list diseases that mosquitoes can transmit in Attica region and got 1 point if they could identify at least one correctly. A fourth question which was initially added to the knowledge score, asked what other animals can get diseases from the mosquitoes. However, since it was less clear how to score various answers regarding risk to domestic animals for the respondent identified pathogens, it was finally decided to exclude this question from the knowledge score.

#### 2. Attitude score

The attitude score was estimated based on answers to four questions aimed at elucidating respondent attitudes concerning exposure to mosquitoes and management activities. Question one asked if and how often respondents were bothered by mosquitoes. With this question, we assumed that respondents who are often bitten by mosquitoes will be favorably predisposed to undertake preventive measures. Thus, respondents who answered, “every day” or “a few days a week” scored 1 while any other answer scored 0 points. The second question asked the respondents if mosquitoes altered their outdoor activities. If they did, the score was 1, otherwise they scored 0 points. The third question of the group, asked the respondents if they undertake any actions to control mosquitoes in their properties. If they answered yes, they scored 1 point otherwise 0 points. The respondents were then asked the kind of actions undertaken, although these activities were not scored. The fourth question asked them how concerned they were by diseases carried by mosquitoes. In a scale from one to five, those ansewered four or five, scored 1 while any other answer scored 0.

### Household investigation

At each household, we surveyed the yard and/or the veranda in case of block flats. For every visit, we recorded the permanent and occasional breeding sites. In the first visit (July 2017), for each household, we kept a detailed file with the type of the water-holding containers present: yard containers, structural containers and trash containers. Following the same procedure in the second visit (November 2017), we updated the initial file. For the households in the “2^nd^ Control Group” (only one visit in November 2017) we followed the same procedure as in July 2017. During all household surveys we recorded the containers found for each container type, including structural, yard (functional containers associated with yard use), and trash (discarded containers).

### Data analysis

A total of 64 questionnaires were distributed 37 were distributed at the time of the initial mosquito survey in July and an additional 27 questionnaires were distributed to the “2^nd^ Control group” at the time of the 2^nd^ household visit in November. The relationships between the parts of the KAP questionnaires are presented in Fig 1.

**Figure 1.**
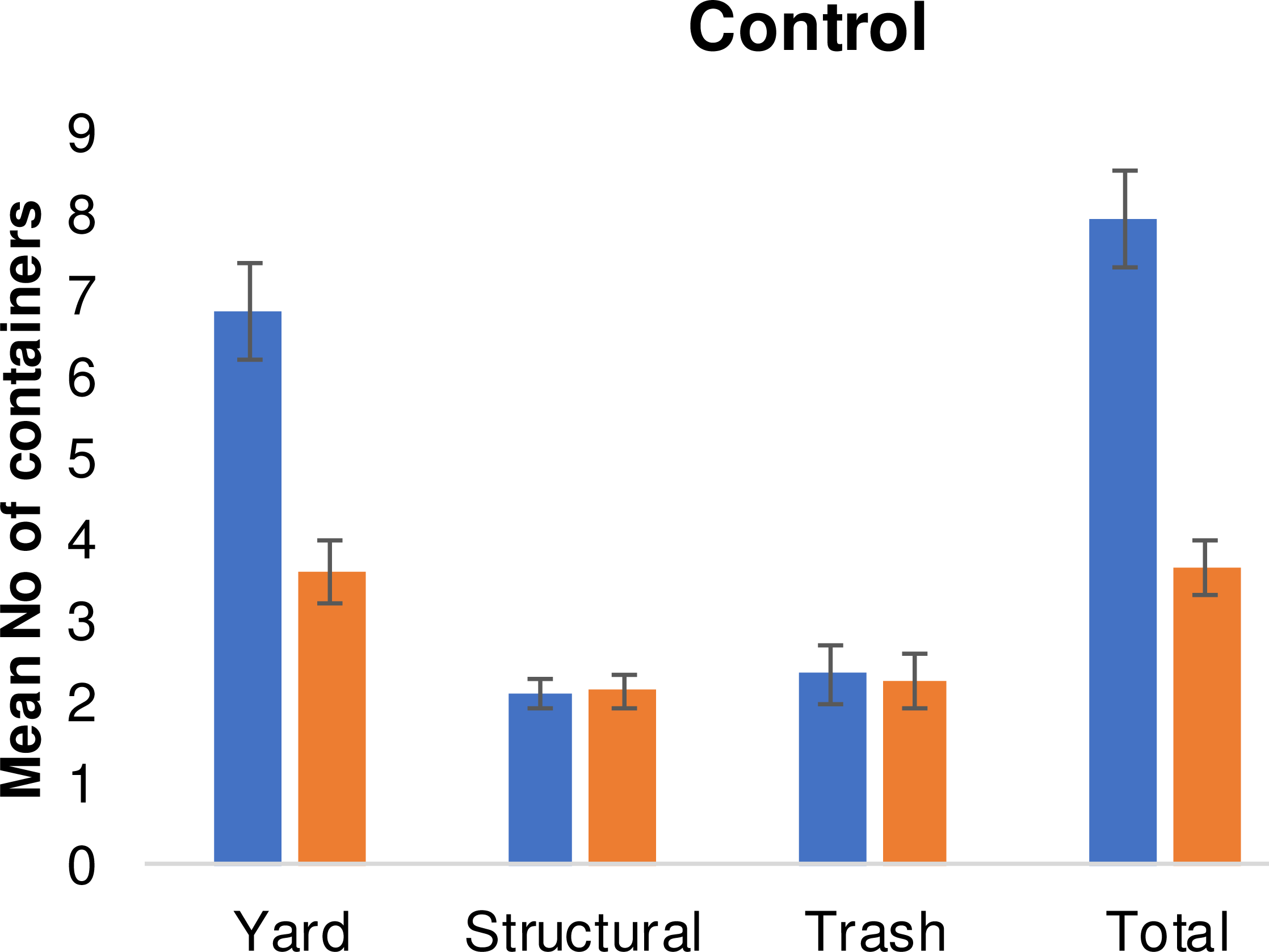
Diagram of the practice followed with KAP questionnaires and educational material for managing mosquito breeding sites

All statistical analyses were completed using the R statistical software (version 3.3.1). Statistical significance was evaluated at alpha=0.05. Analyses focused on answering the following four questions:

**Question 1:** Are there demographic or housing variables that are associated with greater numbers of existing mosquito habitat at the initial survey period?

Welch Two Sample T-tests were used to evaluate the importance of binary predictor variables and to assess pre-intervention differences in container habitat. Pearsons product-moment correlation was used to similarly assess the importance of the ordered, non-binary predictor variables, including education, household income, and year in residence.

**Question 2:** Is the initial container habitat abundance associated with reported resident knowledge, attitude or practice metrics?

The answer to this question was approached through three specific questions (below) about how knowledge, attitude and practices (KAP) responses relate to the abundance and type of source container habitat. For each sub-question, a Pearson’s product-moment correlation was used to evaluate the association between KAP response and observed habitat abundance for each household. Differences in KAP responses among the three treatment groups (education, control, 2^nd^ control) were evaluated using ANOVA.

i. Was the level of resident knowledge (understanding of mosquito ecology) associated with the initial numbers of container habitat? (Knowledge metrics)
ii. Was container number lower if residents were more bothered by mosquitoes or more worried about disease? (Attitude metrics)
iii. Were resident reported practices of avoiding and managing mosquitoes associated with yard-level container habitat? (Practice metrics)

**Question3:** Could demographic, KAP, or housing information predict relative container reductions between the two visits?

A proportional change metric was calculated for each residence sampled twice, as the difference between the second and the initial count divided by the initial count. Pearson’s product-moment correlation was used to evaluate associations between the proportional change in container habitat in a yard and the resident survey response, as well as demographic or housing information, unless the demographic response was binary, when groups were compared using a t test.

**Question 4:** Did an intervention/education treatment result in greater habitat reduction?

A t-test was used to evaluate the effect of the imposed treatment intervention between treatment and control groups. Finally, a generalized linear model (glm function with Poisson error) was used to evaluate differences in container habitat between the treatment group, the original control group, and 2^nd^ control group. Because building administrator presence was an important predictor of both initial container habitat and overall reductions, this predictor variable was further evaluated as part of the generalized linear model.

### Ethics statement

The current study did not involve endangered or protected species and KAP (knowledge, attitudes and practices) study does not reveal confidential information regarding human participants. The ethical committee of Benaki Phytopathological Institute concluded that the current study did not involve human as research subject, thus was not a subject to review.

## Results and Discussion

After the distribution of education material, there was a 54% and 71% total container reduction in the first control group and the treatment group respectively. The reduction was mainly attributed to the reduction of the yard containers which accounted for the 76% and 77% of the total containers in the first control group and the treatment group. The mean number of yard containers in the 1^st^ visit was 6.8 (±4.6) and 9.5 (±7.3) in the control and treatment group respectively (Fig 2). The same values were reduced to 3.6 (±2.5) and 4.5 (±4.0) respectively at the 2^nd^ visit (Fig 3). Relatevely to the structural containers, the mean number in the 1^st^ visit was 2.1 (±1.0) and 1.5 (±0.9) in the control and treatment group respectively. The same values were not reduced at the 2^nd^ visit. The mean number of trash containers in the 1^st^ visit was 2.3 (±1.9) and 2.5 (±2.9) in the control and treatment group respectively. The same values were slightly reduced to • (±1.2) and 2.0 (±4.7) respectively at the 2^nd^ visit. The reduction in the total number of containers was greater among the respondents with higher formal education attainment (Fig 4 and Fig 5). For this group of respondents the average number of total containers was reduced from 8.5 (±5.0) to 6.4 (±5.0) in the first and second visit respectively. Respondents with less formal education reduced the average total number of containers from 9.2 (±6.6) to 8.4(±7.1).

**Figure 2.**
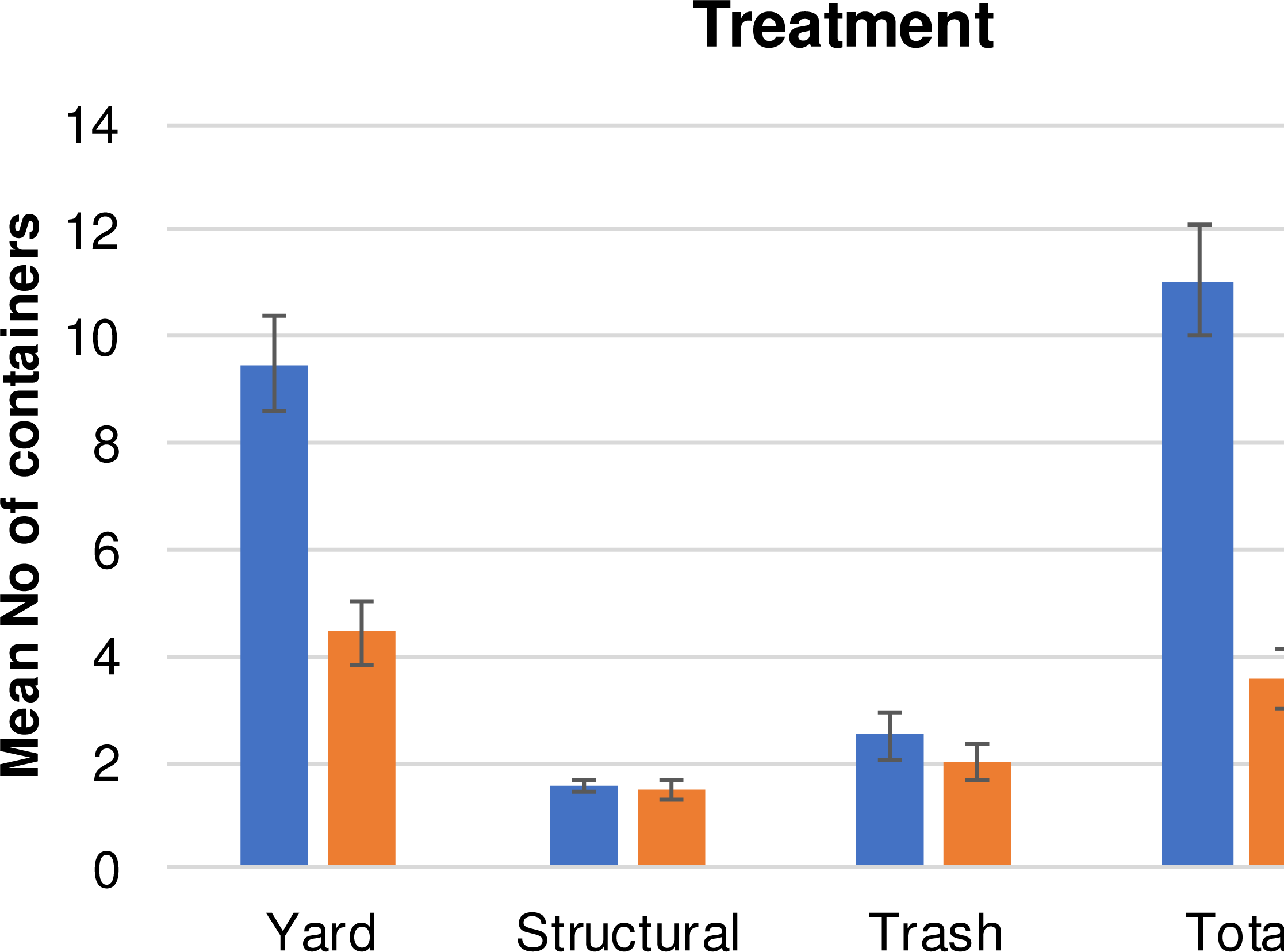
Mean (±SE) number of containers by type of container (yard, structural, trash, total) in the 1^st^ and 2^nd^ visit for the control group.

**Figure 3.**
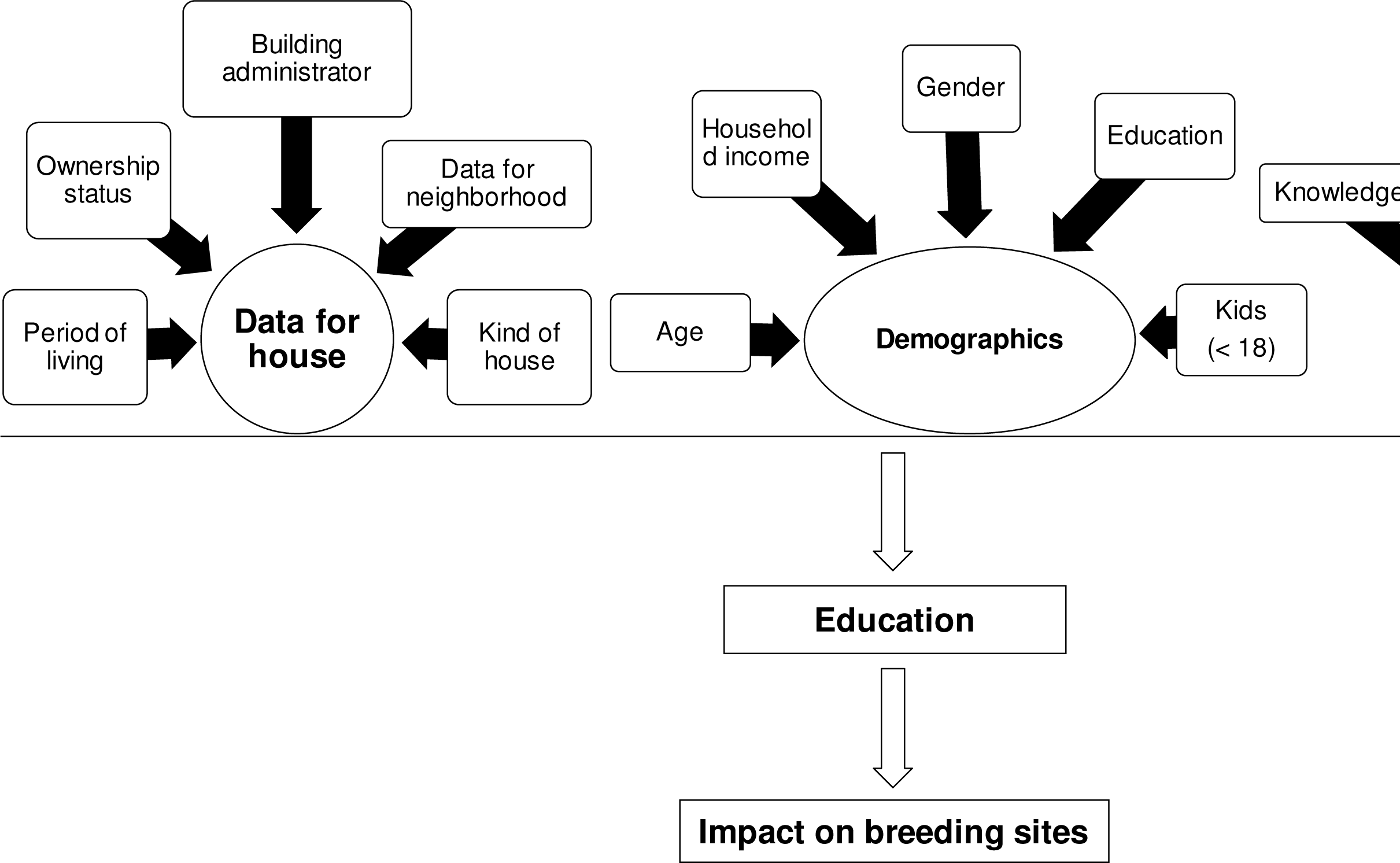
Mean (±SE) number of containers by type of container (yard, structural, trash, total) in the 1^st^ and 2^nd^ visit for the treatment group.

**Figure 4.**
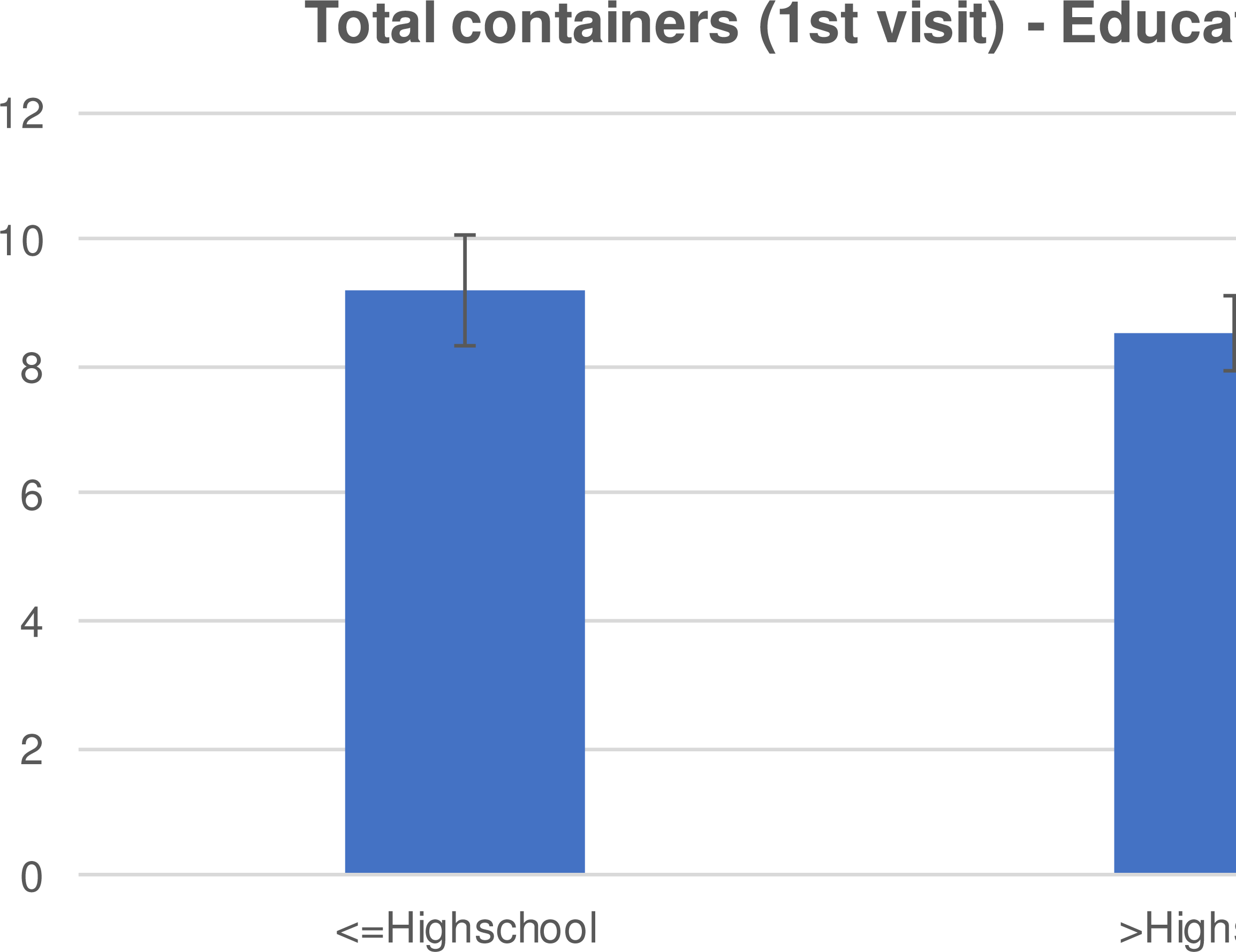
Mean (±SE) total containers during the 1^st^ visit by education level.

**Figure 5.**
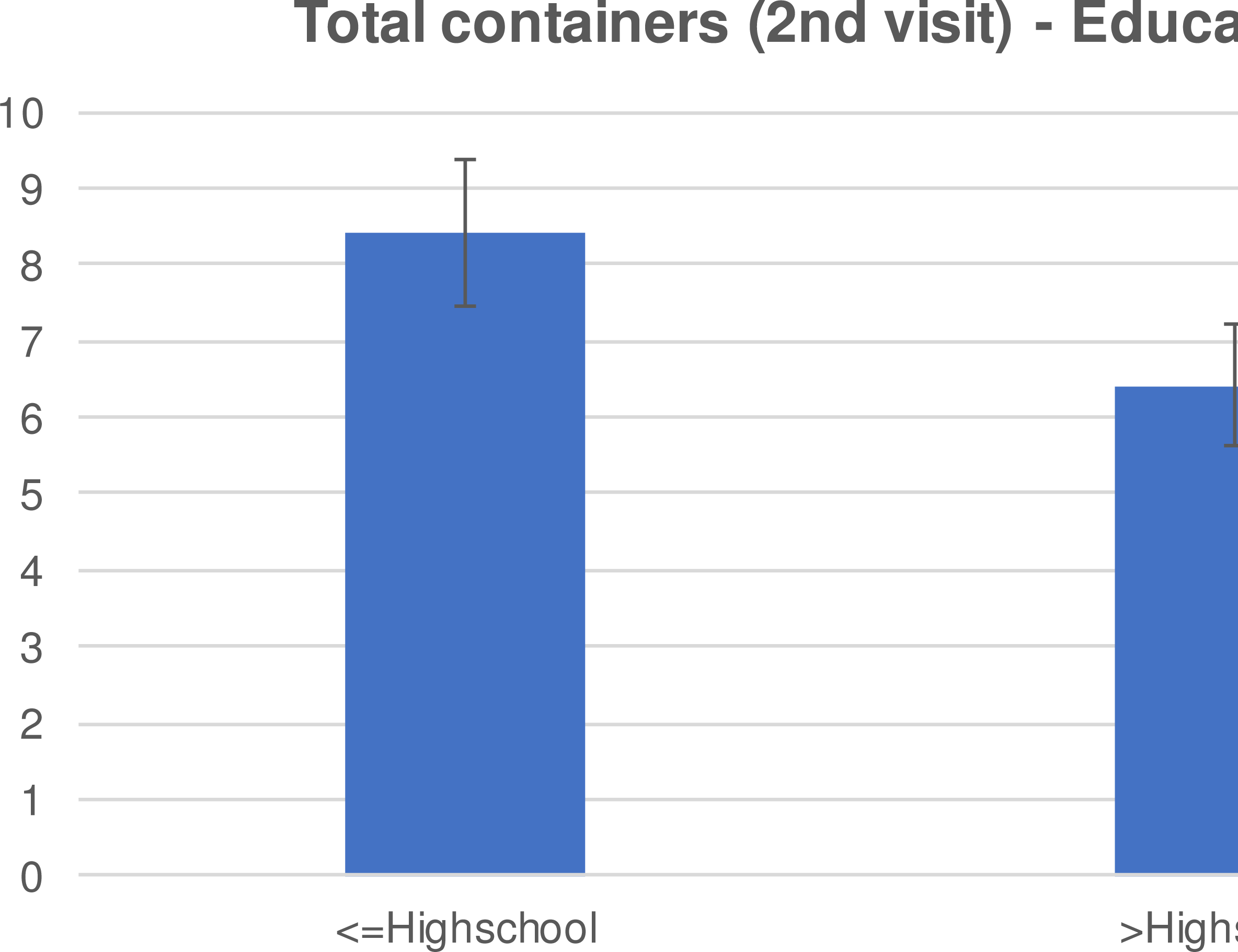
Mean (±SE) total containers during the 2^nd^ visit by education level.

More than 30% of the respondents across all three treatment groups were able to identify the correct diseases transmitted with mosquitoes in the area of Attica (Table 1). The diseases identified were West Nile and Dengue Virus as well as malaria. Malaria received the majority of the responses (21.8%) most probably due to the malaria cases, identified as locally transmitted in 2011 in the Attica region (www.cdc.gov) that resulted in raising awareness on the issue. However, a large percentage of the respondents did not know about the source habitats of the local mosquitoes. Only 18,6% identified “standing water” as important breeding habitat. An important part of the respondents (54%) did consider themselves responsible for mosquito control.

**Table 1.**
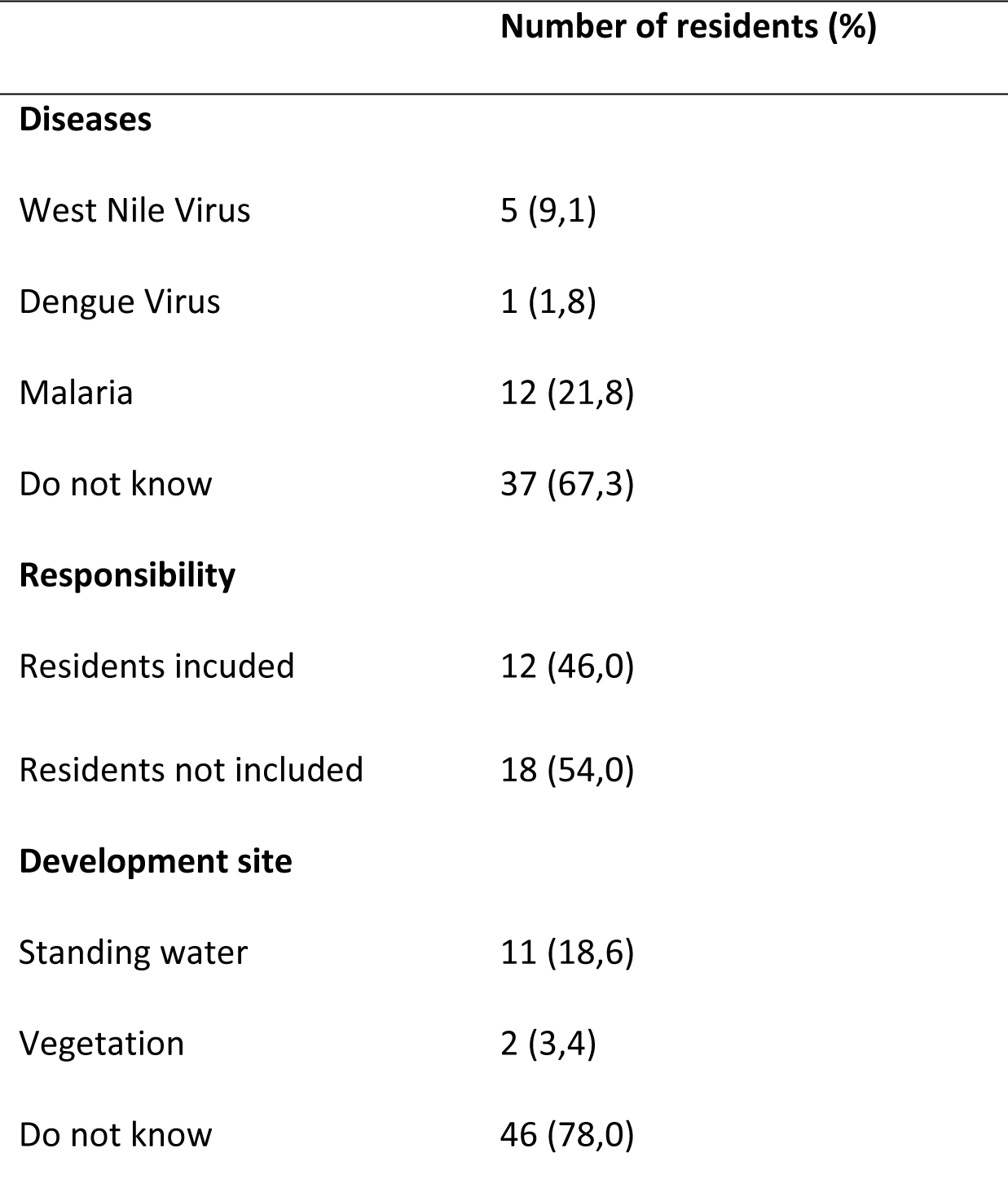
Questionnaire Responses on mosquito knowledge

|  | Number of residents (%) |
| --- | --- |
| **Diseases** |  |
| West Nile Virus | 5 (9,1) |
| Dengue Virus | 1 (1,8) |
| Malaria | 12 (21,8) |
| Do not know | 37 (67,3) |
| **Responsibility** |  |
| Residents incuded | 12 (46,0) |
| Residents not included | 18 (54,0) |
| **Development site** |  |
| Standing water | 11 (18,6) |
| Vegetation | 2 (3,4) |
| Do not know | 46 (78,0) |

***Question 1:*** The statistical analysis showed that there was no significant difference in the initial mean number of containers per yard between the control and treatment groups (control mean: 7.9 ±5.0, treatment mean: 11.0 ± 9.2), or for any subset of container types. Total, Yard or Trash container numbers were not significantly associated with any of the measured demographic or housing variables. However, there were fewer structural containers per yard in buildings with an administrator (t=2.09, d.f.=33, p=0.044) and when the surveyed residence was part of a larger (> 4 apartments) building (t=-3.55, d.f.=33, p=0.001).

***Question 2(i)***-<u>Knowledge approach:</u> A minority of residents knew where mosquitoes breed (18.6%) and only 46% felt that residents had any responsibility for managing breeding habitat (Table 1). The overall mean resident knowledge score was 0.86 (sd=0.85). The knowledge score could range between 0 and 5, with “5” being the “perfect” knowledge score. The knowledge score was not a significant predictor of initial container habitat. The knowledge scores did not differ among the three education treatment groups at the initial visit (F=1.1, df=2 and 61, p >.05) which was made before the actual treatment.

*Question 2(ii)-* <u>Attitude approach</u>: Was container number lower if residents were more bothered by mosquitoes or more worried about disease?

A majority of residents reported (Table 2) being bothered by mosquitoes at least several times a week (88.1%). The “perfect” attitude score equals to 7 while the minimum score is 0. The average level of worry related to diseases carried by mosquitoes was 3.9 (sd 1.3). Resident attitude score was negatively associated with the initial numbers of trash (cor=-0.38, t=-2.45, df=35, p=0.02) and Yard (cor=-0.33, t=-2.03, df=35, p=0.05) but not Structural (cor=-0.06, t=-0.38, df=35, p=0.70) containers. In general, residents attitude scores associated with greater concern had yards with fewer of the removable containers. Attitude scores did not differ among the three education treatment groups at the initial visit (F=0.66, d.f.=2 and 61, p >.05).

**Table 2.**
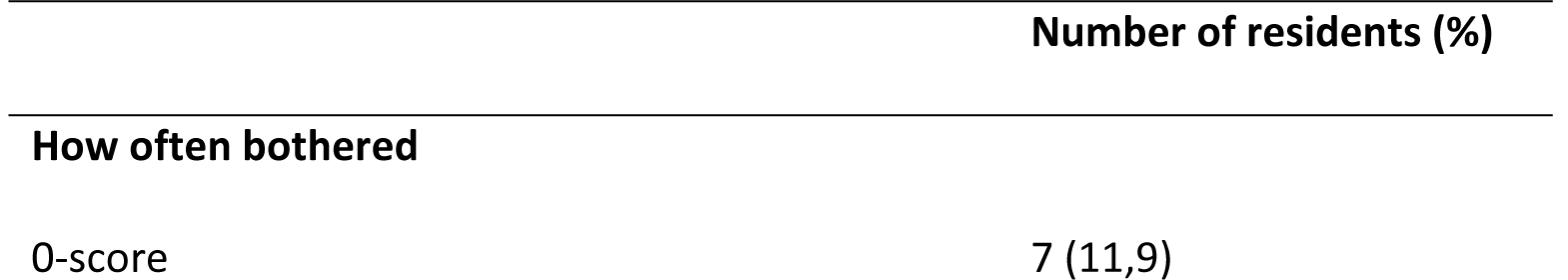
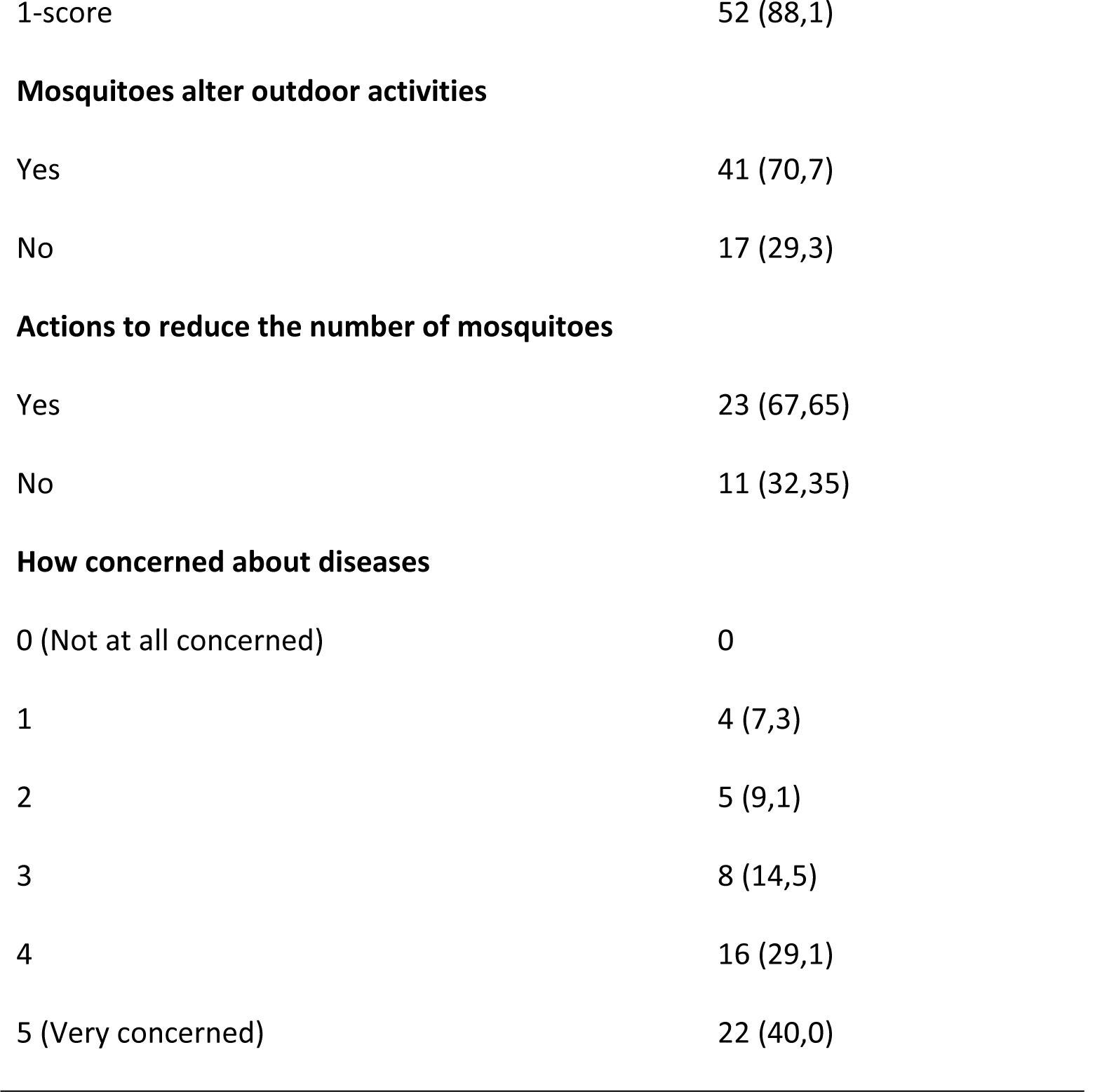
Questionnaire Responses on attitude towards mosquitoes

|  | Number of residents (%) |
| --- | --- |
| **How often bothered** |  |
| 0-score | 7 (11,9) |
| 1-score | 52 (88,1) |
| **Mosquitoes alter outdoor activities** |  |
| Yes | 41 (70,7) |
| No | 17 (29,3) |
| **Actions to reduce the number of mosquitoes** |  |
| Yes | 23 (67,65) |
| No | 11 (32,35) |
| **How concerned about diseases** |  |
| 0 (Not at all concerned) | 0 |
| 1 | 4 (7,3) |
| 2 | 5 (9,1) |
| 3 | 8 (14,5) |
| 4 | 16 (29,1) |
| 5 (Very concerned) | 22 (40,0) |

Respondents who were also building administrators were more motivated to achieve source reduction (Fig 6). The motivation score varied with the ownership status and also with the house owners presenting higher motivation score compared to the tenants (Fig 7). Overall motivation varied with level of formal education attained. Lower formal education level was associated with higher motivation score (Fig 8).

**Figure 6.**
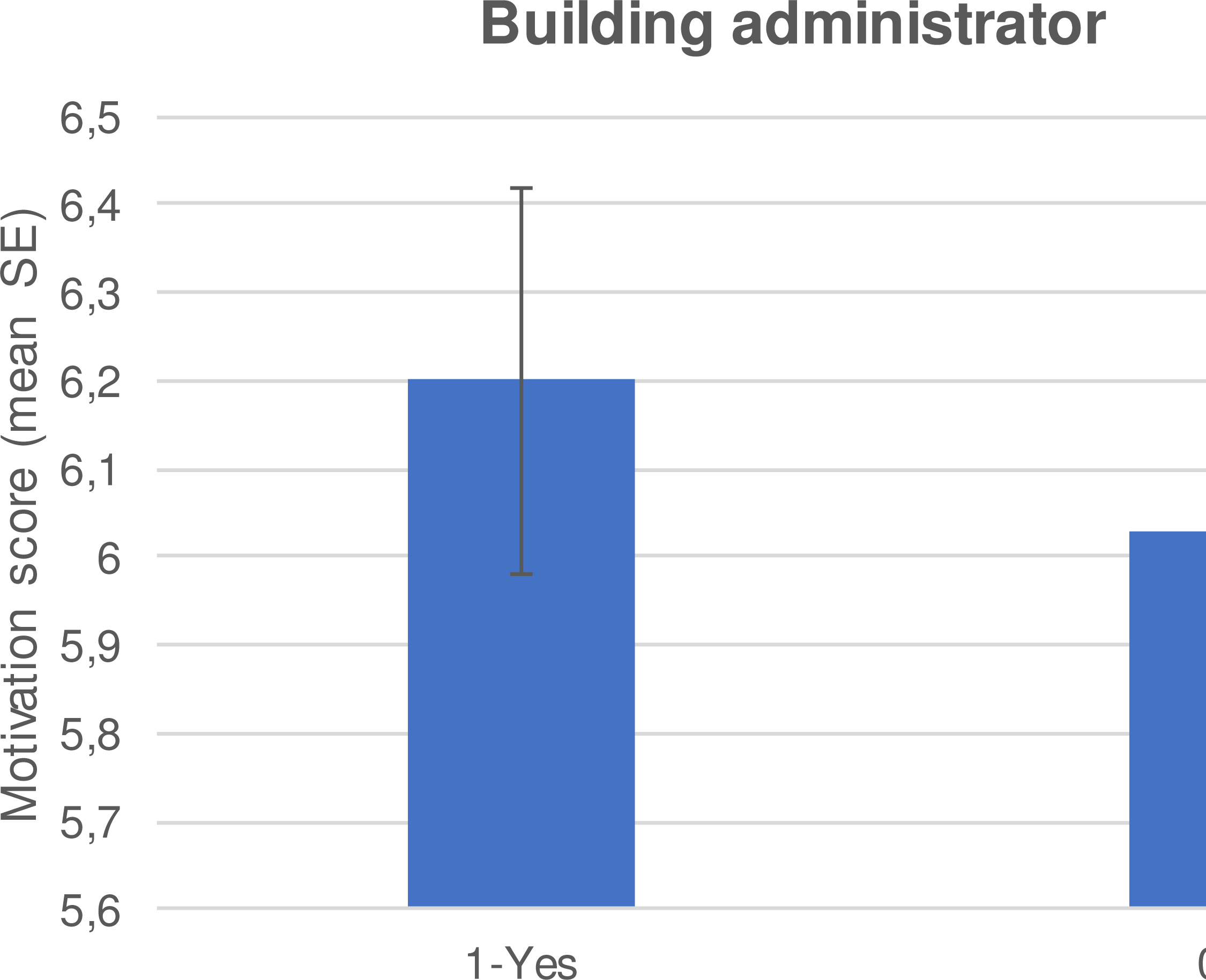
Mean (±SE) motivation score whether the respondent was a building administrator or not.

**Figure 7.**
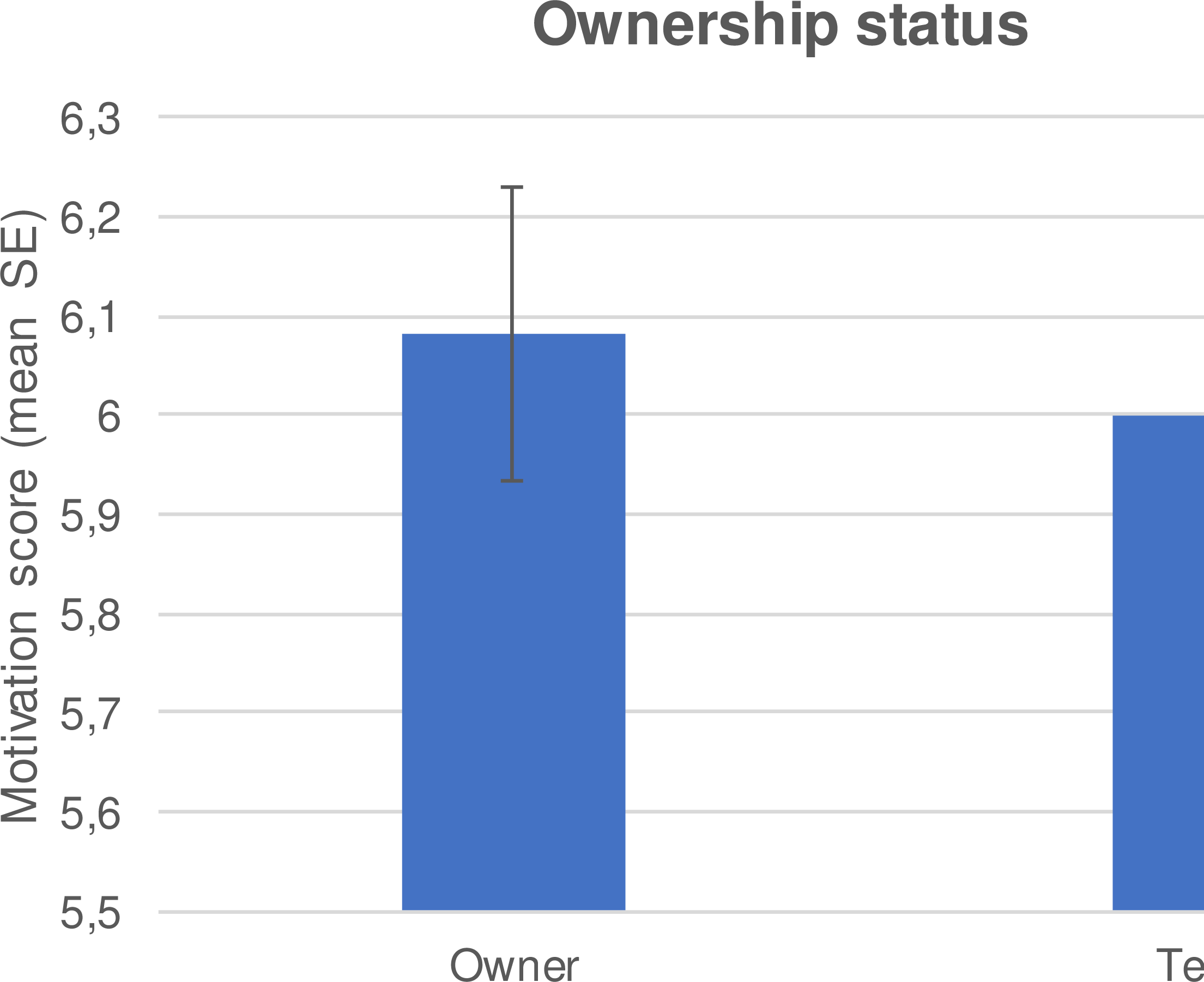
Mean (±SE) motivation score by ownership status.

**Figure 8.**
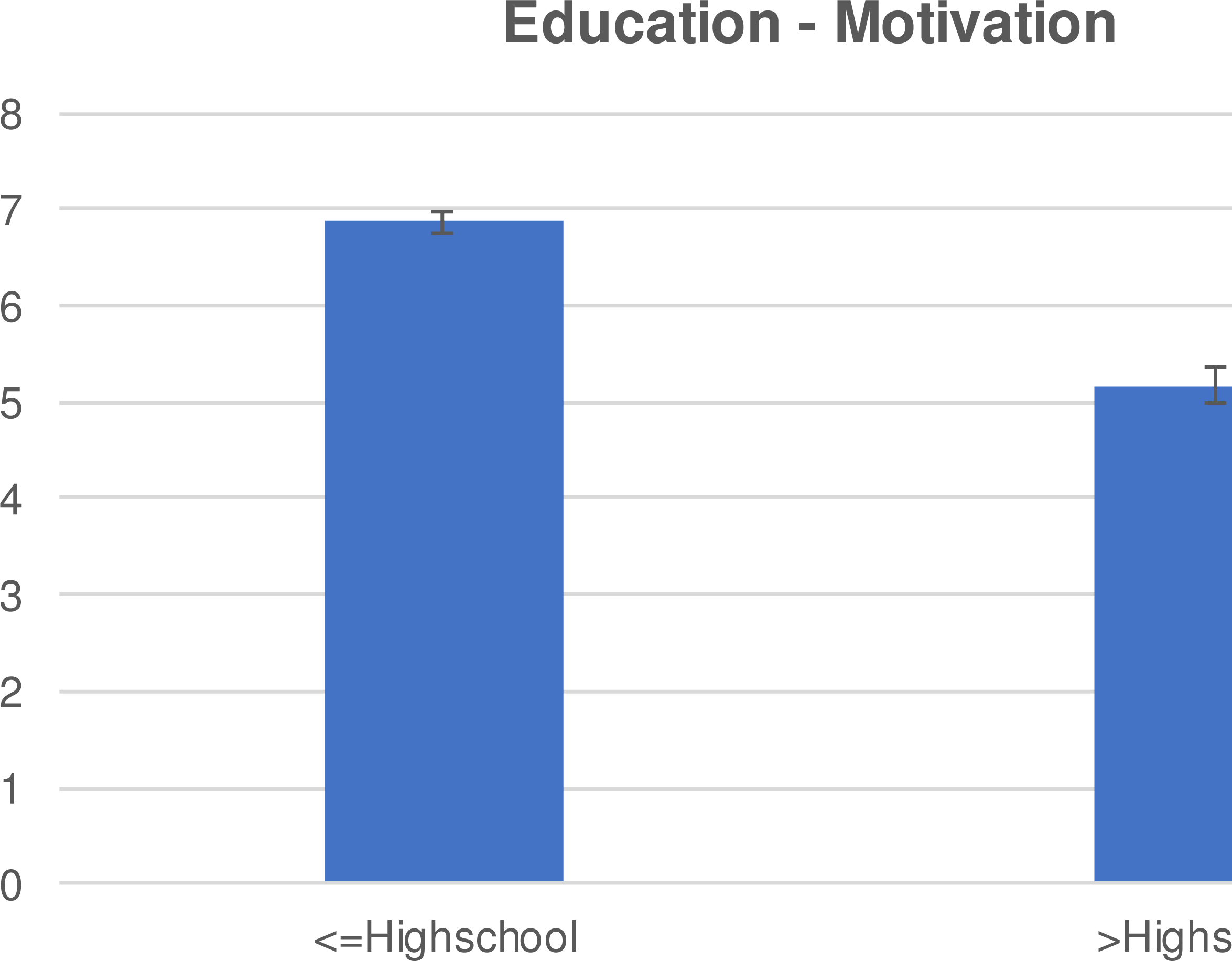
Mean (±SE) motivation score by education level.

Question 2(iii) - A majority of residents reported altering their outdoor activities due to mosquito nuisance (70.7%) and taking some action to limit mosquito exposure (67.7%). In our study, most of the residents reported actions which were related to source reduction, such as emptying water containers, metallic mosquito mesh and application of larvicides.

Although there were no significant associations between reported practices and total numbers of Trash or Structural containers, the numbers of Yard containers were negatively associated with reported avoidance and exposure management (cor=-0.37, t=-2.34, df=35, p=0.02). Practice scores did not differ among the three education treatment groups at the initial visit (F=0.54, df=2 and 61, p >.05).

***Question3-*** There were fewer containers during the second sampling period in all but two households, where container number was unchanged (1 household each in Treatment and Control group). This is reflected in the proportional reductions shown in the Table 3.

**Table 3.**
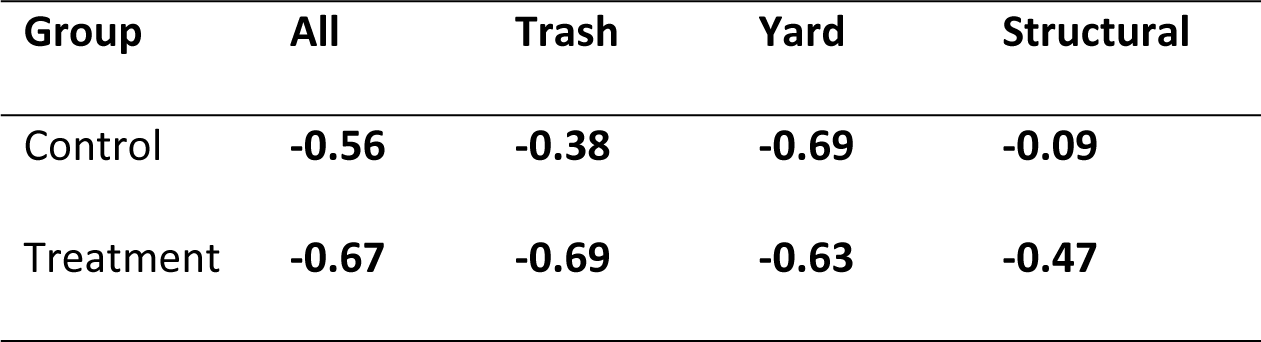
Proportional change in container habitat in control and education treatment groups.

The reduction in total numbers of container habitat was largest when a building administrator was present (t=-2.21, df= 33, p=0.03). The only other significant association was a greater reduction of trash containers in yards with older residents (t=-2.74, df=12, p=-.018). There were no other significant associations between resident demographics (or housing type) and the changes in container habitat between sampling periods. Likewise, there were no significant associations between knowledge, attitude or practice scores and the change in container habitat between sampling periods. Table 4 summarizes the demographic information of the respondents.

**Table 4.** Questionnaire Responses on demographic information

|  | Number of residents (%) |
| --- | --- |
| **Age** |  |
| 18-45 | 25 (41,0) |
| 46-60 | 21 (34,4) |
| >60 | 15 (24,6) |
| **Household income (€)** |  |
| <5000 | 3 (8,8) |
| 5000-10000 | 10 (29,4) |
| 10000-20000 | 14 (41,2) |
| 20000-30000 | 4 (11,8) |
| >30000 | 2 (5,9) |
| Unemployed | 1 (2,9) |
| **Ownership status** |  |
| Owner | 43 (74,1) |
| Tenant | 15 (25,9) |
| **Education level** |  |
| Less than Highschool | 6 (11,1) |
| Highschool | 20 (37,0) |
| College/BSc | 21 (38,9) |
| MSc/PhD | 7 (13,0) |
| **Type of house** |  |
| 4 or less apartments | 19 (30,6) |
| More than 4 apartments | 43 (69,4) |
| **Building administrator** |  |
| Yes | 23 (37,1) |
| No | 39 (62,9) |

***Question 4-*** There were no significant differences in Total, Yard or Trash container habitat reductions between the treatment and original control groups. There was a marginally greater reduction in structural container habitat in the treatment group (t=1.90, df=18, p=0.07).

There were significantly fewer container habitats in the second survey of yards receiving the education treatment when compared to the 2^nd^ control yards (estimate= 1.30, z=9.59, p=<0.01) but there was no significant difference between treatment and the original control (z=10.15, p=0.89). There was no significant interaction between Treatment group and having a building administrator. The education treatment group had significantly fewer numbers of both yard and structural container habitats in the second survey when compared to the 2^nd^ control (structural estimate= 2.27, z=6.583, p<0.001; yard estimate= 1.14, z=7.24, p<0.001) but not between treatment and the original control (structural; z=1.68, p=0.09; yard; z=-1.14, p=0.25). Education treatment effect was not significantly associated with presence of a building administrator. There were no significant differences in the numbers of trash habitat containers between treatment and the 2^nd^ control (z=-1.52, p=0.12) or control (z=0.36, p=0.72) groups.

One of the major goals of our study was to test whether the use of an educational intervention could enhance the community engagement and result in source reduction. This is the first time that KAP study is implemented in Greece and considered important for preventing mosquito bites and consequently to reduce the risk of mosquito-borne diseases. In Greece there are currently only imported cases of Chikungunya fever [20] but in areas at risk of entry or establishment and risk for disease transmission resident education could play important role [21].

Since this was the first time in Greece that a KAP study was implemented, we included a 2^nd^ control group during the second visit-inspection (November 2017) aiming to evaluate behaviors that may be influenced because of our first inspection (first visit on July 2017). Results revealed a visit bias between treatment and control group and even though the reduction of total containers was important, our education campaign found to have no effect on the habitat reduction. After including the 2^nd^ control group, the effect of the printed educational material was found significant between the treatment group and the 2^nd^ control group. The above results strongly suggest that only the presence of scientific staff inspecting possible habitats in their properties, could be enough to stimulate practices towards source reduction. The presence of mosquito personnel counting containers might have raised concern about being fined for producing mosquitoes in their property [17]. Their concern was further amplified since they did not receive any educational material.

The changes in any types of containers were not significantly associated with the motivation metrics. Nevertheless, residents attitude scores associated with greater concern, had yards with fewer of the removable containers. This was probably because of the greater level of nuisance which urged them to limit their exposure to mosquitoes and to the larger number of breeding sites.

Our study also found that the total number of containers found in the first visit did not differ between respondents with different formal education level. However, at the second visit, and after the distribution of the educational material, the decrease in the number of total containers was higher among respondents with higher formal educational attainment. This result suggests that educational interventions need to be better designed in order to make greater gains when targeted to households with less formal education. The abovementioned results are particularly interesting when combined with respondents from households with higher educational level were considered less motivated to take action to control mosquitoes in their properties, still they seem to have accomplished greater source reduction in their households. The explanation for this may be their greater experience in receiving, elaborating and finally adopting education messages which made them more prone to adopt the control measures suggested by the educational material.

## Conclusion

Findings from our study suggest that a single visit-inspection of a trained mosquito expert in the households and the inspection of the potential breeding sites in their yards can influence the residents’ behavior towards source reduction. However, educational interventions alone with printed education material cannot enhance significant community participation and source reduction and funds may be better used to support other strategies. As proposed by [22] intersectoral co-operation with the involvement of local health services, trained vector control personnel, civil authorities and the community could contribute to converting information into practice.

## Acknowledgments

The authors would like to thank the deputy mayor of the Municipality, Mrs V. Andrikopoulou and Mrs A. Fikiri, Mrs A. Georgopoulou and all the staff from the Environmental Department (Environment Unit and the Green) of Palaio Faliro for their active help and participation in the distribution and collection of the questionnaires and identify households to participate in the study.

